# Test-retest reproducibility of *in vivo* oscillating gradient and microscopic anisotropy diffusion MRI in mice at 9.4 Tesla

**DOI:** 10.1101/2021.08.04.455122

**Authors:** Naila Rahman, Kathy Xu, Mohammad Omer, Matthew D. Budde, Arthur Brown, Corey A. Baron

## Abstract

**Background and Purpose:** Microstructure imaging with advanced diffusion MRI (dMRI) techniques have shown increased sensitivity and specificity to microstructural changes in various disease and injury models. Oscillating gradient spin echo (OGSE) dMRI, implemented by varying the oscillating gradient frequency, and microscopic anisotropy (µA) dMRI, implemented via tensor valued diffusion encoding, may provide additional insight by increasing sensitivity to smaller spatial scales and disentangling fiber orientation dispersion from true microstructural changes, respectively. The aims of this study were to characterize the test-retest reproducibility of *in vivo* OGSE and µA dMRI metrics in the mouse brain at 9.4 Tesla and provide estimates of required sample sizes for future investigations.

**Methods:** Eight adult C57Bl/6 mice were scanned twice (5 days apart). Each imaging session consisted of multifrequency OGSE and µA dMRI protocols. Metrics investigated included µA, isotropic and anisotropic kurtosis, and the diffusion dispersion rate (Λ), which explores the power-law frequency dependence of mean diffusivity. The dMRI metric maps were analyzed with mean region-of-interest (ROI) and whole brain voxel-wise analysis. Bland-Altman plots and coefficients of variation (CV) were used to assess the reproducibility of OGSE and µA metrics. Furthermore, we estimated sample sizes required to detect a variety of effect sizes.

**Results:** Bland-Altman plots showed negligible biases between test and retest sessions. ROI-based CVs revealed high reproducibility for both µA (CVs < 8 %) and Λ (CVs < 15 %). Voxel-wise CV maps revealed high reproducibility for µA (CVs ∼ 10 %), but low reproducibility for OGSE metrics (CVs ∼ 50 %).

**Conclusion:** Most of the µA dMRI metrics are reproducible in both ROI-based and voxel-wise analysis, while the OGSE dMRI metrics are only reproducible in ROI-based analysis. µA and Λ may provide sensitivity to subtle microstructural changes (4 - 8 %) with feasible sample sizes (10 – 15).

## INTRODUCTION

Diffusion MRI (dMRI) provides a non-invasive means to capture microstructure changes in the brain during development, aging, disease, and injury by probing the diffusion of water molecules [1]. The most widely used dMRI techniques are diffusion tensor imaging (DTI) and diffusion kurtosis imaging (DKI). DTI assumes the dMRI signal is entirely characterized by Gaussian diffusion [2] and utilizes a diffusion tensor model to estimate metrics such as mean diffusivity (MD) and fractional anisotropy (FA). DKI provides more information about the underlying tissue via the diffusion kurtosis, which quantifies the deviation from Gaussian diffusion [3]. However, both DTI and DKI are unable to distinguish between microstructural changes and neuron fiber orientation dispersion [2,4], reducing their specificity to microstructural changes in brain regions with crossing fibers. Furthermore, DKI cannot differentiate between different sources of kurtosis (non-Gaussian diffusion) [3].

Probing microstructure with diffusion-weighted sequences beyond the conventional Stejskal-Tanner pulsed gradient spin echo (PGSE) sequence [5], used in DTI and DKI, is currently of broad interest. The aims of these emerging dMRI sequences are to overcome the limitations of DTI and DKI and improve sensitivity and specificity to microstructural changes. In the present work, the reproducibility of *in vivo* oscillating gradient and microscopic anisotropy dMRI, both of which have unique features that go beyond the PGSE sequence, is investigated in mice at 9.4 Tesla.

The conventional PGSE sequence consists of a pair of pulsed gradients applied along a single direction. Here, the diffusion measurement reflects information about diffusion along a single direction and at a single relatively long diffusion time, which is the time allowed for water molecules to probe the local environment. Given hardware constraints, the minimum diffusion times achievable with PGSE sensitize the signal to length scales of tens of micrometers, which is larger than typical axon sizes (∼ 2 µm) and cell sizes (∼ 10 – 30 µm) [6].

To overcome the diffusion time limitations of PGSE, the oscillating gradient spin echo (OGSE) method was developed to modify sensitivity to cellular length scales [7]. OGSE allows different microstructure length scales to be probed by varying the frequency of the oscillating diffusion gradients, which is inversely related to diffusion time. For increasing diffusion times (lower oscillating gradient frequencies), the molecules travel greater distances and interact with more barriers such as cell membranes, resulting in lower observed MD values [8]. As MD is different at the various frequencies, this provides the ΔMD - the metric of interest in OGSE dMRI, the difference in MD between the highest and lowest frequencies applied. By acquiring diffusion data at multiple frequencies, the power law relationship between MD and frequency (f) can be explored via the “diffusion dispersion rate”, Λ [9,10]. Evidence of a linear dependence of MD on the square root of frequency has been demonstrated in healthy and globally ischemic rodent brain tissue [11] and healthy human white matter [12]. Thus, Λ can be calculated as

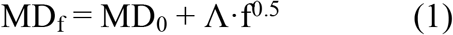

where MD_f_ is the OGSE MD at a frequency f and MD_0_ is the MD at f = 0 [9,10,12]. Since OGSE is sensitive to structural disorder along one dimension [9], changes in the number and morphology of neurite varicosities will result in changes to Λ [10], which potentially makes OGSE an invaluable tool to probe microstructural changes, such as axonal beading, *in vivo* [13,14].

In contrast to the widely used fractional anisotropy metric (FA), which confounds true microstructural changes with fiber orientation dispersion [2], the microscopic anisotropy (µA) metric quantifies water diffusion anisotropy independent of orientation dispersion [15]. To disentangle orientation dispersion from true microstructure changes, the shape of the b-tensor, which describes the strength of diffusion weighting along each direction, is varied via tensor-valued diffusion encoding [16]. Most tensor-valued encoding protocols are based on double diffusion encoding (DDE) techniques or a combination of linear tensor encoding (LTE) and spherical tensor encoding (STE). As DDE sequences are implemented via two consecutive diffusion encoding pulses separated by a mixing time, they require longer TEs than standard LTE/STE sequences to achieve equal b-values [17]. Conventional DTI and DKI utilize only LTE, in which all gradients are along the same axis, so that diffusion is encoded along a single direction at a time. STE, in which the gradients are distributed throughout all directions, sensitizes the signal to diffusion along all directions at the same time. Here, a combination of LTE and STE is utilized to implement microscopic anisotropy (µA) dMRI [4,15]. The µA metric is defined based on the difference in signal between LTE and STE dMRI acquisitions [15,18]. As the LTE signal depends on variance of both isotropic and anisotropic diffusivity, while the STE signal depends only on variance of isotropic diffusivity, diffusional kurtosis estimated from the µA protocol can be separated into two components: anisotropic kurtosis (KLTE – arising from the LTE acquisitions) and isotropic kurtosis (KSTE – arising from the STE acquisitions). Thus, KLTE is a measure of the dispersion in the orientation of diffusion tensors and KSTE is a measure of the variance in the magnitude of diffusion tensors or the mean diffusivity, which can be related to cell size heterogeneity [4].

OGSE and µA dMRI have recently been gaining attention in various disease and injury models and their feasibility has been shown in both preclinical and clinical settings. Importantly, OGSE dMRI can provide measures of mean cell size [19,20] and axonal diameter [21,22], while µA dMRI can provide estimates of cell shape [4]. The OGSE ΔMD metric has shown increased sensitivity, compared to MD alone, in the assessment of hypoxia-ischemia [23] and radiation therapy treatment response [24] in rodents, and in various pathologies in humans, including muscle contraction abnormalities [25], high- and low-grade brain tumor differentiation [26], and neonatal hypoxic-ischemic encephalopathy [27]. Notably, OGSE has helped to identify neurite beading as a mechanism for dMRI contrast after ischemic stroke [13,14]. Preliminary studies in humans have found that µA provides better sensitivity than the conventional FA in distinguishing between different types of brain tumours [4], assessment of multiple sclerosis lesions [28,29], and detecting white matter microstructure changes associated with HIV infection [30]. Furthermore, Westin et al. reported that KLTE and KSTE showed significant differences between controls and schizophrenia patients, while conventional mean kurtosis showed no difference [31]. The feasibility of µA dMRI has been demonstrated in rodents both *in vivo* [32,33] and *ex vivo* [34–36]. *In vivo* preclinical rodent µA studies, which have included predominantly DDE techniques and more recently combined LTE/STE techniques, have shown that KSTE may be particularly sensitive to deep gray matter lesions [37], µA dMRI can enable robust estimation of microscopic diffusion kurtosis (or intra-compartmental kurtosis) [38], and promising results from a rodent model of epilepsy indicating that microscopic diffusion kurtosis can provide improved characterization of tissue microstructure changes, compared to conventional DKI [39].

As dMRI has reached the forefront of tissue microstructure imaging [40], there is a need to establish the reproducibility of these emerging methods. While the reproducibility of DTI and DKI has been investigated extensively [41–44], to the best of our knowledge, no test-retest assessment of OGSE and µA dMRI has been done at an ultra-high field strength. The aim of this work was to assess test-retest reproducibility of *in vivo* OGSE and µA dMRI in mice at 9.4 Tesla and provide estimates of required sample sizes, which is essential in planning future preclinical neuroimaging studies involving models of disease/injury.

## METHODS

### Subjects

All animal procedures were approved by the University of Western Ontario Animal Use Subcommittee and were consistent with guidelines established by the Canadian Council on Animal Care. Eight adult C57Bl/6 mice (four male and four female) were scanned twice 5 days apart. The sample size was chosen to reflect common practice in pre-clinical imaging studies [23,39,45]. Before scanning, anesthesia was induced by placing the animals in an induction chamber with 4 % isoflurane and an oxygen flow rate of 1.5 L/min. Following induction, isoflurane was maintained during the imaging session at 1.8 % with an oxygen flow rate of 1.5 L/min through a custom-built nose cone. The mouse head was fixed in place using ear bars and a bite bar to prevent head motion.

### *In vivo* MRI

*In vivo* MRI experiments were performed on a 9.4 Tesla (T) Bruker small animal scanner equipped with a gradient coil set of 1 T/m strength (slew rate = 4100 T/m/s). A single channel transceive surface coil (20 mm x 25 mm), built in-house, was fixed in place directly above the mouse head to maximize signal-to-noise ratio (SNR). Each dMRI protocol was acquired with single-shot spin echo echo-planar-imaging (EPI) readout with scan parameters: TR = 10 s; in-plane resolution = 175×200 µm; slice thickness = 500 µm; 30 slices to acquire the full brain; field-of-view = 19.2 × 14.4 mm^2^; partial Fourier imaging in the phase encode direction with 80% of k-space being sampled; 45 minutes scan time. For each dMRI protocol, a single reverse phase encoded b = 0 s/mm^2^ volume was acquired at the end of the diffusion sequence for subsequent use in TOPUP [46] and EDDY [47] to correct for susceptibility and eddy current induced distortions. Averages were acquired separately on the scanner and combined using in-house MATLAB code to correct for frequency and signal drift associated with gradient coil heating [48]. Anatomical images were also acquired for each subject within each session using a 2D T2-weighted TurboRARE pulse sequence (150 μm in-plane resolution; 500 μm slice thickness; TE/TR = 40/5000 ms; 16 averages; total acquisition time = 22 min).

#### Oscillating Gradient Spin Echo (OGSE) dMRI

OGSE dMRI was performed with five oscillating gradient frequencies of 0 Hz, 50 Hz, 100 Hz, 145 Hz, and 190 Hz, as shown in **Figure 1** (A – E). The 0 Hz frequency refers to the conventional PGSE sequence. The frequencies were chosen based on a hypoxic-ischemic injury study in mice [23], where the frequencies ranged from 0-200 Hz, which enables probing length scales between 1.2 – 4.2 µm. Other scan parameters included: gradient duration = 11 ms; gradient separation = 5.5 ms; TE = 39.2 ms; 5 averages; b = 800 s/mm^2^; 10 diffusion encoding directions. 10 b = 0 s/mm^2^ volumes were interspersed evenly throughout the acquisition. 10 diffusion encoding directions chosen here combined the 4 (tetrahedral) [12] or 6 direction encoding schemes [45,49] commonly used in OGSE.

**Fig 1.**
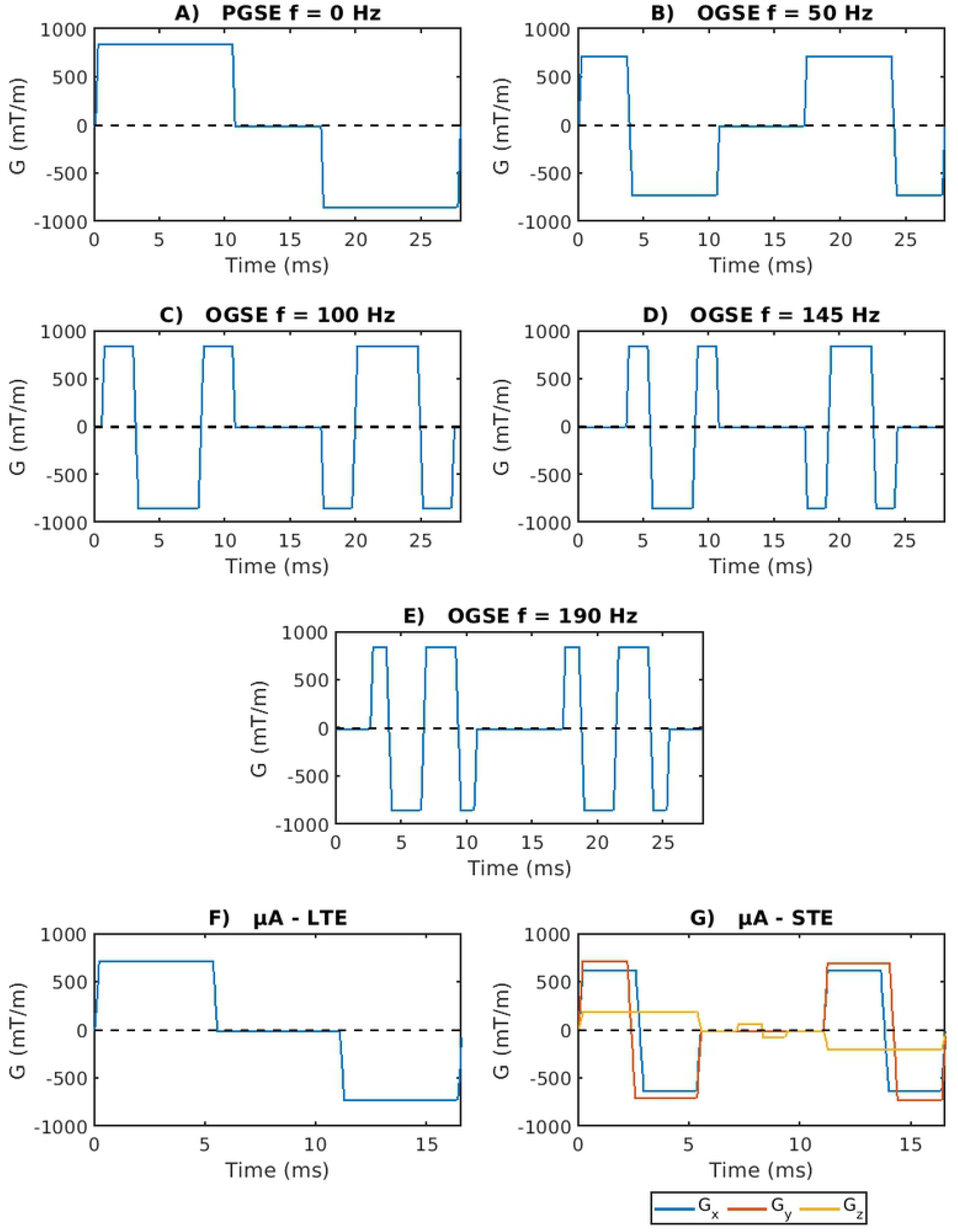
Schematic representations of the gradient waveforms used for the OGSE and µA protocols. Diffusion encoding blocks have been inserted on both sides of a 180° pulse and implicit gradient reversal due to the 180° pulse has been applied. **A – E** show the sequences used in the OGSE protocol, which include one PGSE waveform (**A**) and 4 OGSE waveforms with gradient oscillation frequencies of 50 Hz, 100 Hz, 145 Hz, and 190 Hz (**B – E**). **F** and **G** show the LTE and STE waveforms respectively, used in the µA protocol.

#### Microscopic Anisotropy (µA) dMRI

The STE dMRI gradient waveforms implemented here were similar to the protocol in Arezza *et al*. [18]. The μA sequence was implemented with linear (LTE) and spherical tensor (STE) encodings, as shown in **Figure 1** (F – G), at b = 2000 s/mm^2^ (30 directions for each of LTE and STE) and b = 1000 s/mm^2^ (12 directions). Other scan parameters included: gradient duration = 5 ms; gradient separation = 5.54 ms; TE = 26.8 ms; 3 averages. 8 b = 0 s/mm^2^ volumes were interspersed evenly throughout the acquisition.

### Image Processing

Images were pre-processed using PCA denoising [50] and Gibbs ringing correction from the MRtrix3 package [51], followed by TOPUP [46] and EDDY [47] from FMRIB Software Library (FSL, Oxford, UK) [52]. Brain masks were produced using the skull stripping tool from BrainSuite (v. 19b) [53]. Image registration was performed using affine and symmetric diffeomorphic transforms with ANTs software (https://github.com/ANTsX/ANTs) [54]. Region-of-interest (ROI) masks were acquired from the labeled Allen Mouse Brain Atlas [55]. Since registration to an atlas is time-consuming, only one anatomical T2-weighted scan was chosen (the “chosen T2”) to be registered to the atlas. All other anatomical T2-weighted images were registered to the chosen T2. Non-diffusion weighted (b0) volumes were registered to the corresponding anatomical images (from the same subject at the same timepoint). All dMRI volumes were registered to the corresponding anatomical space using the transforms resulting from the previous step (b0 → corresponding T2). For ROI-based analysis, the inverse transforms resulting from these two registration steps (corresponding T2 → chosen T2 → atlas) were then used to bring the labeled atlas to the corresponding T2 space for each subject at each timepoint. Binary masks for each ROI were generated by thresholding the labeled atlas. Each mask was eroded by one voxel, except for the corpus callosum masks, to minimize partial volume errors within a given ROI. The binary masks were visually inspected to ensure good registration quality. Furthermore, to perform whole brain voxel-wise analysis of all subjects across both timepoints, all dMRI volumes were registered to the chosen T2 space using transforms from two registration steps (b0 → corresponding T2 → chosen T2). For voxel-wise analysis targeted to specific ROIs, the labeled atlas was registered to the chosen T2 space.

From the OGSE data, maps of MD at each frequency were generated using MRtrix3 [51,56]. ΔMD was calculated as the difference between MD acquired at the highest frequency (190 Hz) and MD acquired at the lowest frequency (0 Hz). To characterize the power law relationship between MD and OGSE frequency (f) [9], the slope of linear regression of MD with f^0.5^, the diffusion dispersion rate (Λ), was calculated. From the µA data, maps of µA, K_LTE_, K_STE_, and microscopic fractional anisotropy (µFA), which is the normalized counterpart of µA, were generated using an optimized linear regression technique based on the diffusion kurtosis model, described by Arezza *et al*. [18].

### Data Analysis

The test-retest dataset is available online [57]. Measurement reproducibility was explored for both ROI-based analysis and whole brain voxel-wise analysis, since both are common analyses techniques in neuroimaging. To mitigate partial volume errors from cerebrospinal fluid (CSF), voxels with MD (0 Hz) > 0.9 µm^2^/ms were omitted from the analyses of all scalar maps. The ROI analysis focused on five different tissue regions: corpus callosum, internal capsule, hippocampus, cortex, and thalamus. Bland-Altman analysis was performed for both ROI-based and voxel-wise analysis to identify any biases between test and retest measurements. For both analysis techniques, the scan-rescan reproducibility was characterized using the coefficient of variation (CV). The CV reflects both the reproducibility and variability of these metrics and allows calculation of the sample sizes necessary to detect various effect sizes. CVs were calculated between subjects and within subjects to quantify the between subject and within subject reproducibility respectively. The between subject CV was calculated separately for the test and retest timepoints as the standard deviation divided by the mean value across subjects 1– 8. These two CV values were then averaged for the mean between subject CV. The within subject CV was calculated separately for each subject as the standard deviation divided by the mean of the test and retest scans. The 8 within subject CVs were then averaged to determine the mean within subject CV. Following the procedure presented in van Belle [58], the between subject CVs, from the ROI analysis, were used to determine the sample size required per group to detect a defined biological effect between subjects in each ROI. Assuming paired t-tests, the standard deviations of the differences between test-retest mean values across subjects, were used to determine the sample size required to detect a defined biological effect within subjects in each ROI [59]. The minimum sample sizes, using the between and within subject approaches, were both determined at a 95 % significance level (α = 0.05) and power of 80 % (1−β = 0.80).

#### ROI Analysis

The mean MD was calculated for each ROI at each frequency. For each ROI, ΔMD was calculated as the difference between the mean MD at 190 Hz and the mean MD at 0 Hz. The apparent diffusion dispersion rate, Λ, was determined for each ROI by performing a least square fit of the mean MD (in each ROI) to f^0.5^. Metrics from the µA protocol were averaged for each ROI after voxel-wise fitting. Bland-Altman and CV analyses were performed using the mean values.

#### Voxel-wise Analysis

ΔMD maps were generated by subtracting the MD maps at 0 Hz from the MD maps at 190 Hz. Λ maps were generated by performing a least square fit of MD to f^0.5^ for each voxel. Voxel-wise Bland-Altman and CV analyses were performed for each metric using the scalar maps (ΔMD, Λ, and scalar maps from the µA protocol).

## RESULTS

Representative parameter maps are shown in **Figure 2**. MD (190 Hz) has an overall higher intensity than MD (0 Hz). ΔMD shows selective enhancement of distinct regions in the brain - the dentate gyrus (part of the hippocampal formation) is shown with white arrows. As expected, ΔMD and Λ show similar contrast. ROI-based fitting of Λ showed the expected trends with f^0.5^ in all ROIs and at both test and retest time-points (**Figure 3**). The µA and µFA maps also show similar contrast. K_LTE_ highlights white matter structures as expected and K_STE_ is homogenous throughout the brain, although very high in CSF regions.

**Fig 2.**
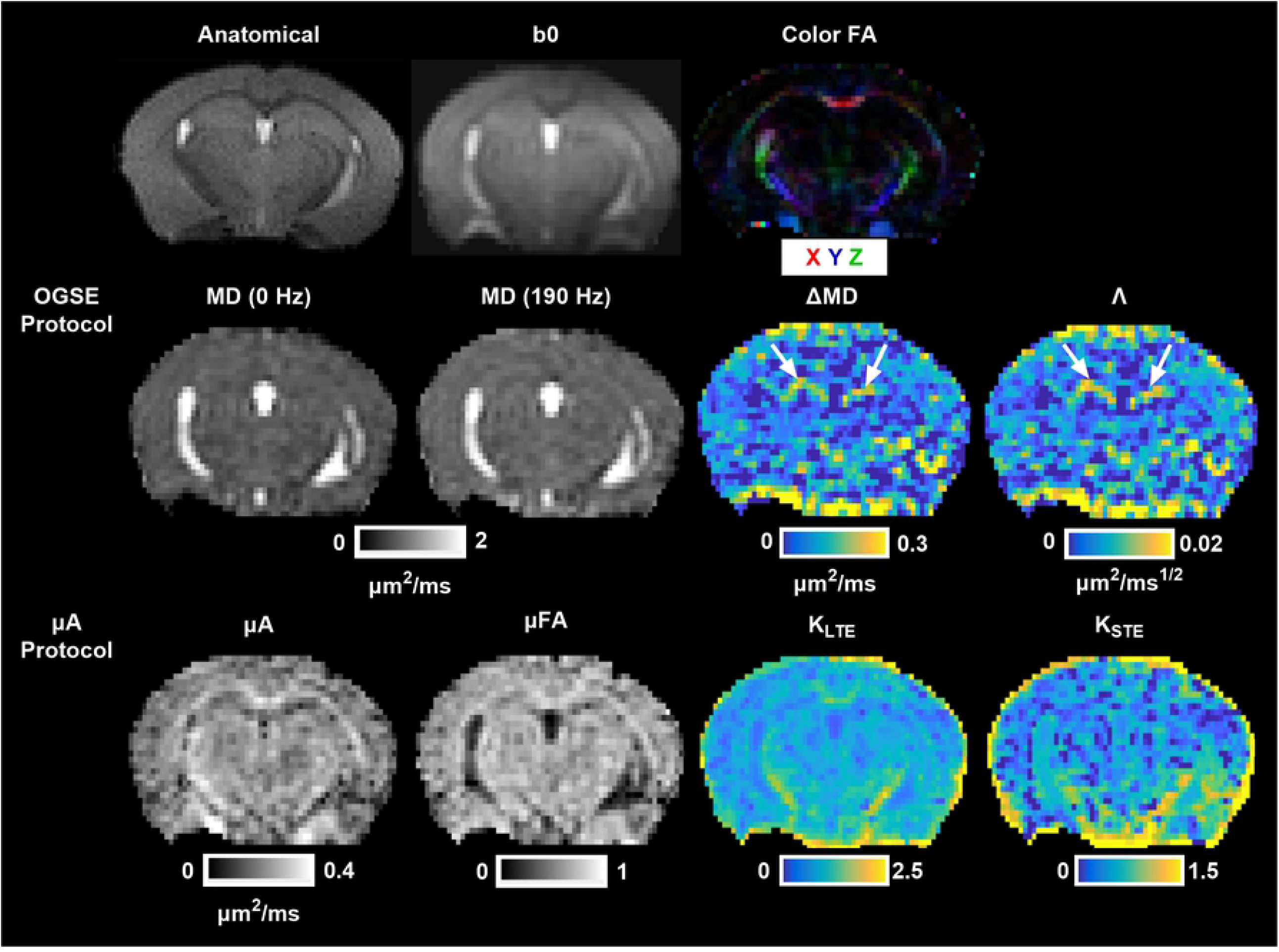
Example axial cross sections from a single subject showing an anatomical T2-weighted image, a non-diffusion weighted image (b0), and a color fractional anisotropy map (Color FA), where the colors represent the primary direction of diffusion. Parameter maps from the OGSE protocol (MD (0 Hz): Mean Diffusivity at 0 Hz; MD (190 Hz): Mean Diffusivity at 190 Hz; ΔMD: the difference between MD (190 Hz) and MD (0 Hz); Λ: the apparent diffusion dispersion rate) and the µA protocol (µA: Microscopic Anisotropy; µFA: Microscopic Fractional Anisotropy; K_LTE_: Anisotropic Kurtosis (from linear tensor encodings); K_STE_: Isotropic Kurtosis (from spherical tensor encodings)) are shown. The white arrows in the ΔMD and Λ maps indicate high OGSE contrast in the dentate gyrus.

**Fig 3.**
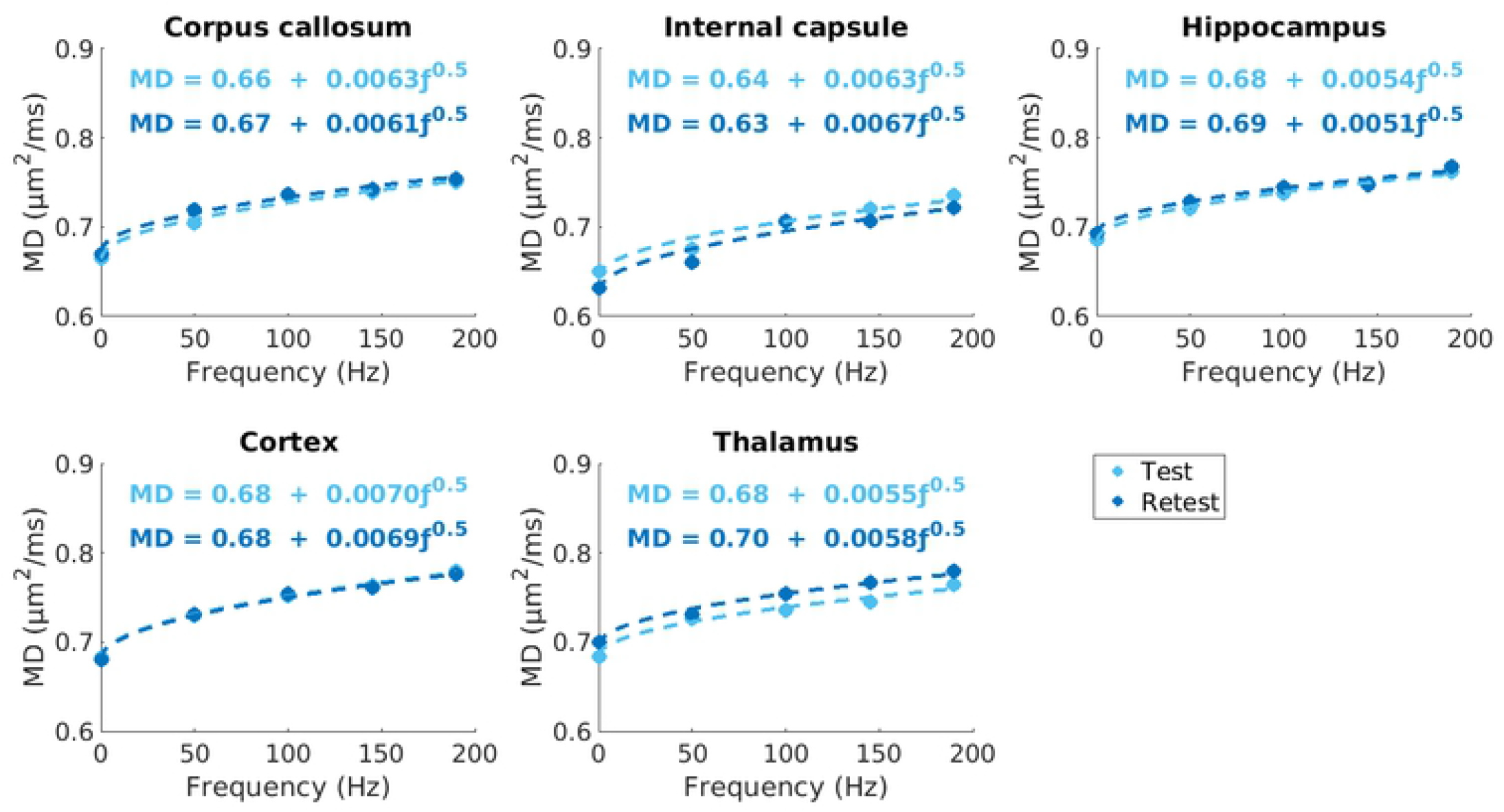
Least square fitting of mean MD values to f^0.5^, depicted by the dotted lines, in each ROI for test and retest timepoints in one mouse. The diffusion dispersion rate, Λ, ranged from 0.0051 – 0.0070 µm^2^/ms^1/2^, depending on the ROI.

### ROI Analysis

Violin plots depict the distribution of the mean values for each metric within each ROI for the eight subjects (**Figure 4**). Across all metrics, the median and interquartile range are similar for test and retest conditions. In general, the smaller ROIs (the internal capsule and the thalamus) show greater distributions, while the larger ROIs (i.e., the cortex) showed much tighter distributions. Bland-Altman plots (**Figure 5**) revealed negligible biases between repeat measurements across all metrics. Lower mean between subject CVs were observed in Λ (3 – 4 %) compared to ΔMD (7 – 18 %), while the within subject CVs were very similar for both metrics, ranging from 3 – 14 % (**Figure 6**). µA and µFA show low between and within subject CVs for all ROIs (ranging from 3 – 8 %), with µFA showing slightly lower CVs. K_LTE_ exhibited much lower between and within subject CVs (4 – 10 %) compared to K_STE_ (10 – 32 %).

**Fig 4.**
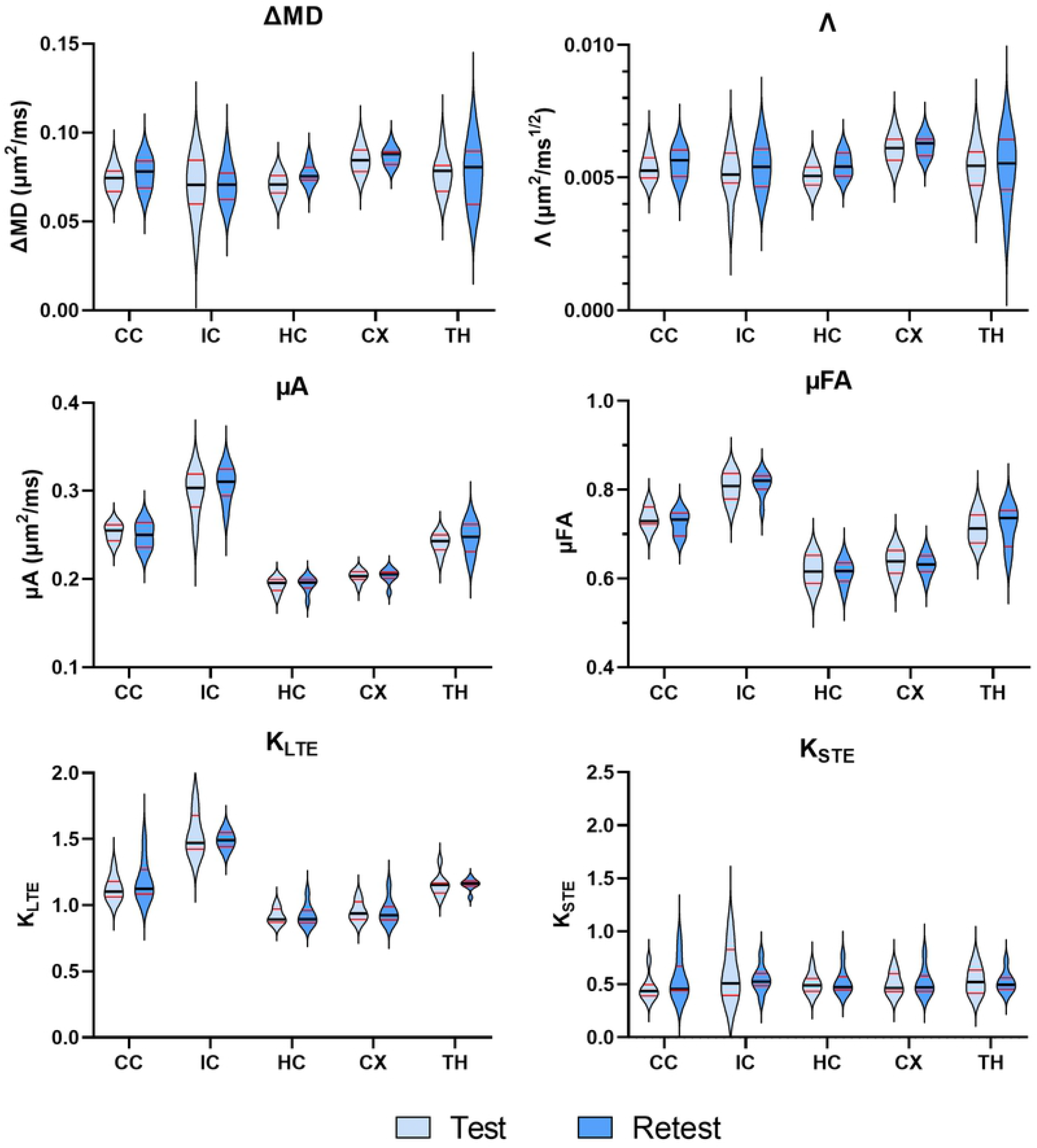
Violin plots showing the distribution of the OGSE metrics (ΔMD and Λ) and the µA metrics (µA, µFA, KLIN, and KST) at the test and retest timepoints (five days apart) for eight subjects in several brain regions. The dark black line represents the median and the red lines depict the interquartile range (25th to 75th percentile). The violin plots extend to the minimum and maximum values of each metric. ROIs are abbreviated as follows: CC – corpus callosum; IC – internal capsule; HC – hippocampus; CX – cortex; TH – thalamus.

**Fig 5.**
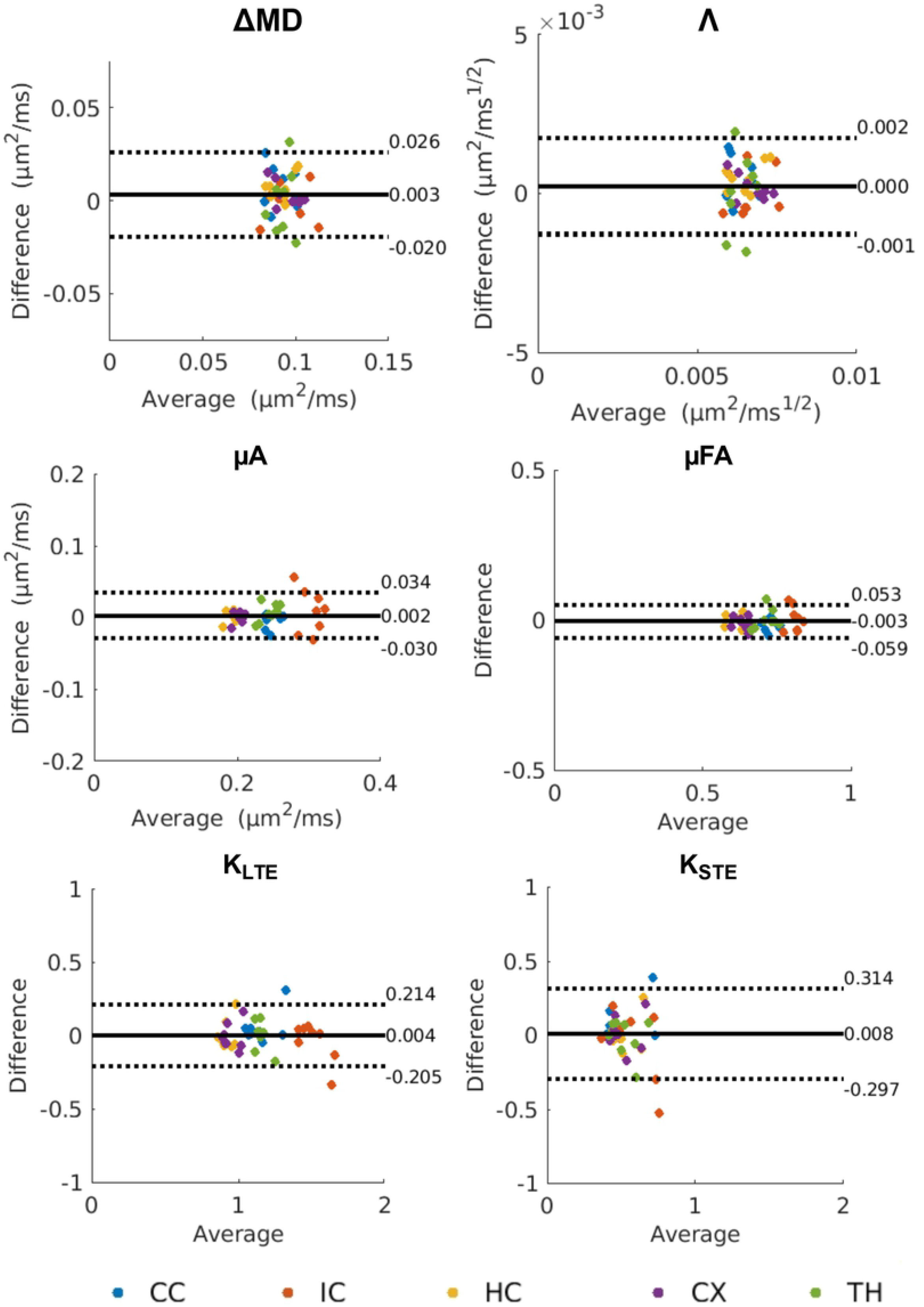
Bland-Altman plots depicting biases between test and retest scans for mean values of OGSE and µA metrics (from the ROI-based analysis). The solid black lines represent the mean bias, and the dotted black lines represent the ±1.96 standard deviation lines. The average of the test and retest mean values is plotted along the x-axis and the difference between the test and retest mean values is plotted along the y-axis. ROIs in the legend are abbreviated as follows: CC – corpus callosum; IC – internal capsule; HC – hippocampus; CX – cortex; TH – thalamus.

**Fig 6.**
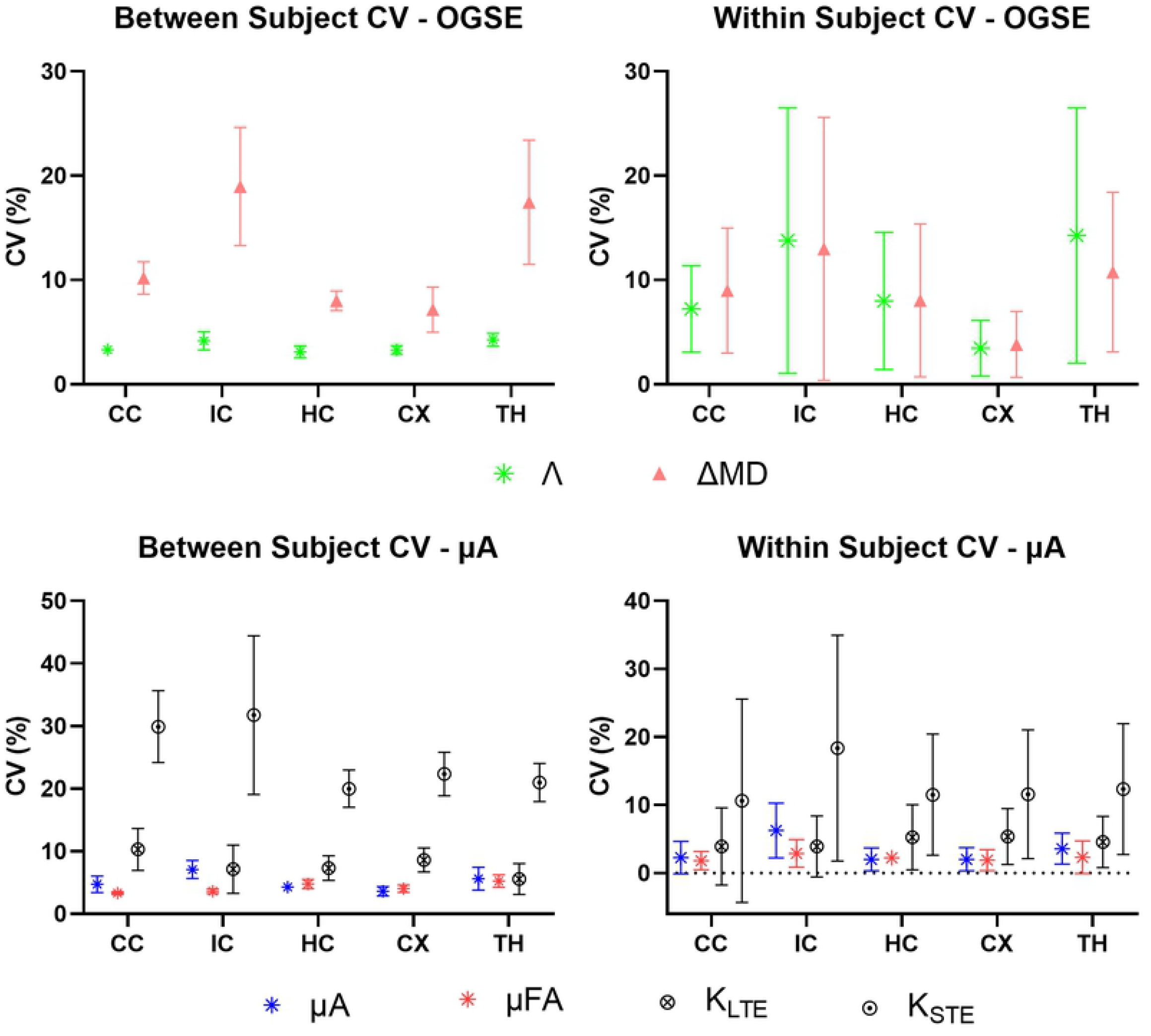
Mean between subject and within subject coefficients of variation (CV) for OGSE and µA metrics for each ROI. Values for the between subject condition represent the mean ± standard deviation over subjects (averaged over the test and retest timepoints). Values for the within subject condition represent the mean ± standard deviation between test and retest (averaged over the eight subjects). ROIs are abbreviated as follows: CC – corpus callosum; IC – internal capsule; HC – hippocampus; CX – cortex; TH – thalamus.

### Voxel-wise Analysis

Bland-Altman plots comparing whole brain test and retest voxels for all eight subjects revealed negligible biases for all metrics (**Figure 7**). However, ΔMD, Λ, and K_STE_ showed greater fluctuations around the estimated bias. The CV maps (**Figure 8**) show very high CVs in the CSF regions (except for the K_STE_ CV maps). Histograms (**Figure 9**) show ΔMD and Λ have the same distribution. Overall, the between and within subject CVs are comparable for all metrics. µA, µFA, and K_LTE_ have comparable CVs with peaks at 10, 8, and 16 % respectively. ΔMD, Λ, and K_STE_ peak around 50 % and have very wide distributions. Whole brain histograms and histograms for specific ROIs (**Supplemental Figure 1**) show similar trends.

**Fig 7.**
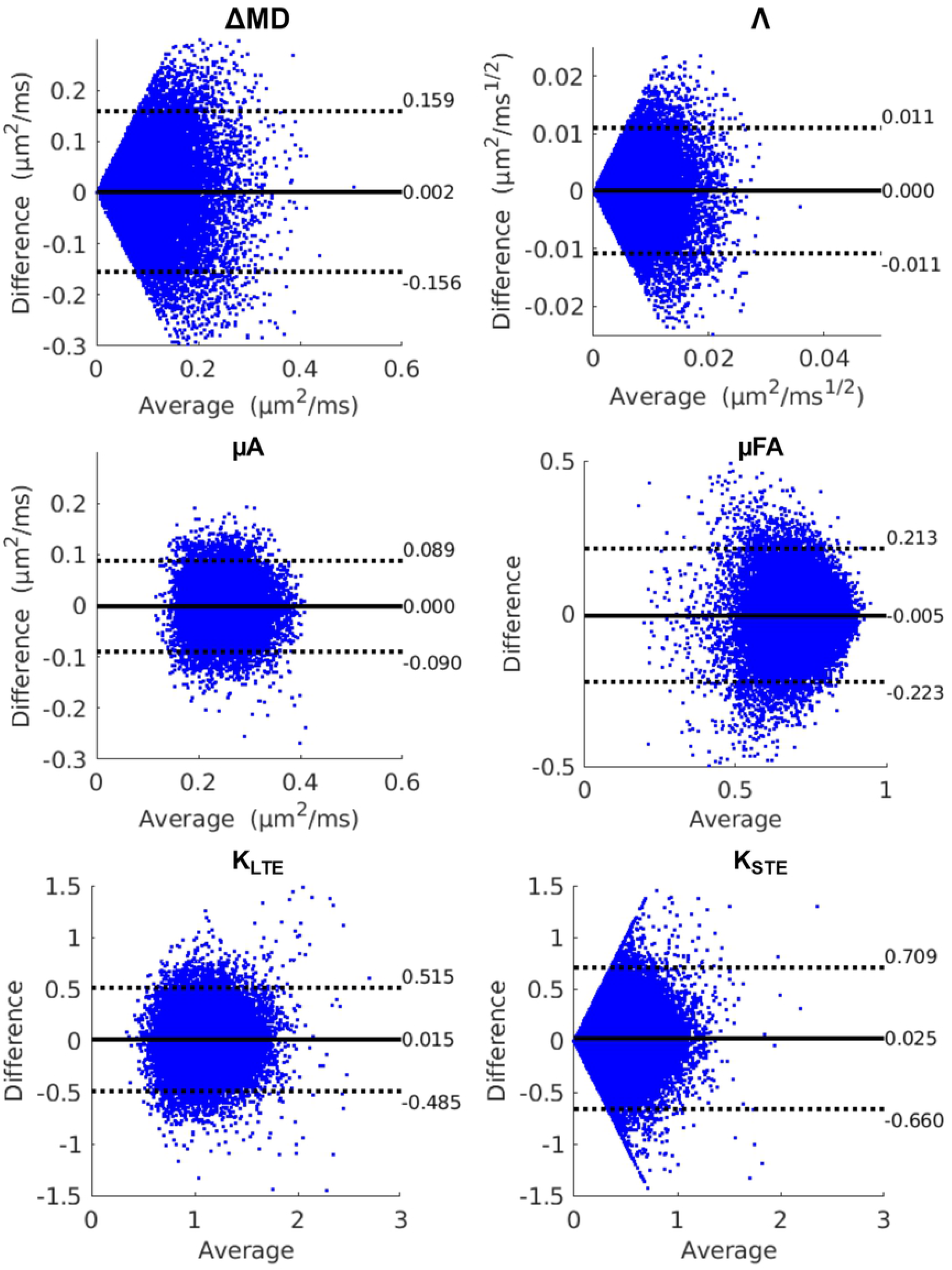
Bland-Altman plots depicting biases between test and retest scans for OGSE and µA metrics from the whole-brain voxelwise analysis for all subjects. The solid black lines represent the mean bias, and the dotted black lines represent the ±1.96 standard deviation lines. The average of the test and retest voxels is plotted along the x-axis and the difference between the test and retest voxels is plotted along the y-axis.

**Fig 8.**
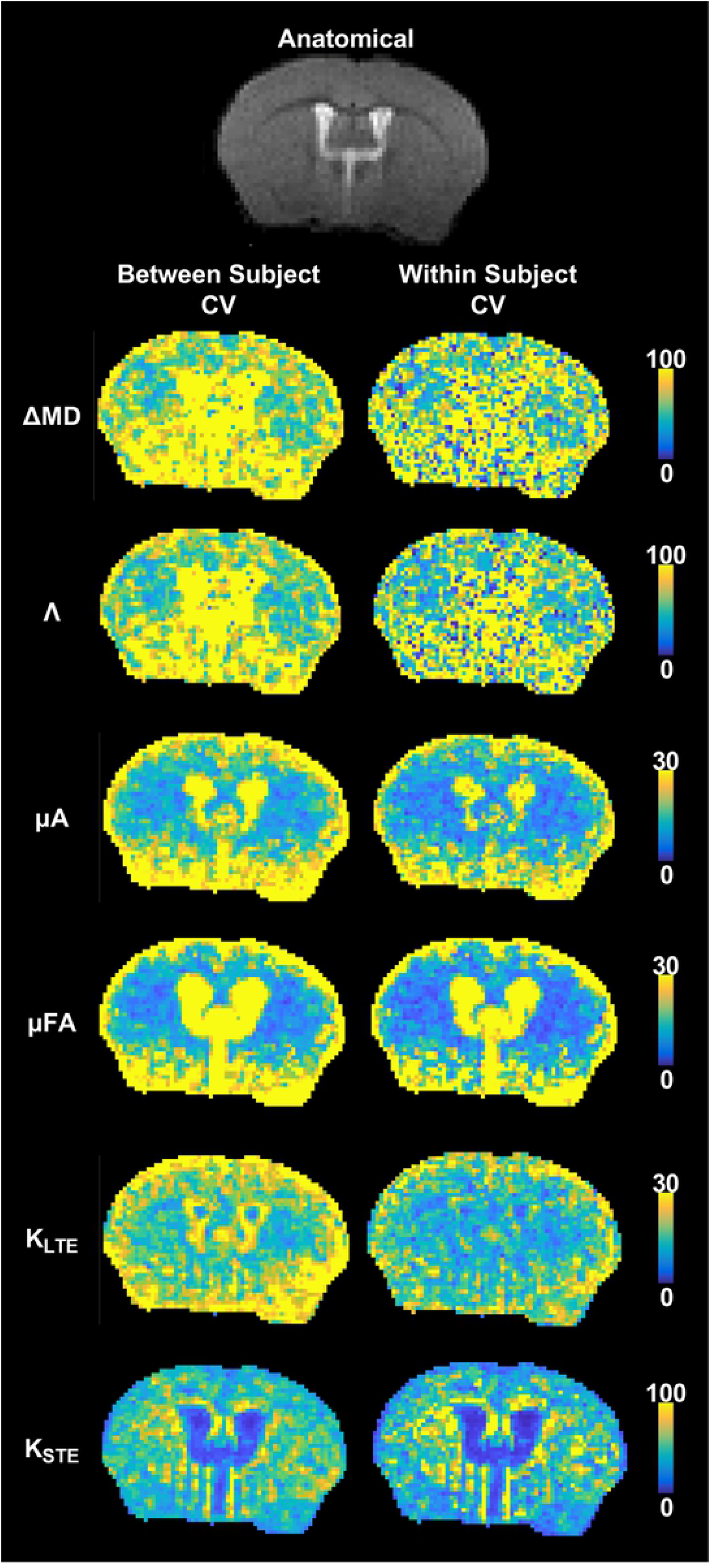
Whole brain average between subject and within subject CV maps. All diffusion data was registered to a single anatomical T2-weighted dataset (representative axial slice shown). Values for the between subject condition represent the mean CV within each voxel averaged over the test and retest timepoints. Values for the within subject condition represent the mean CV within each voxel averaged over all eight subjects. Note that the color bar scale varies between the maps.

**Fig 9.**
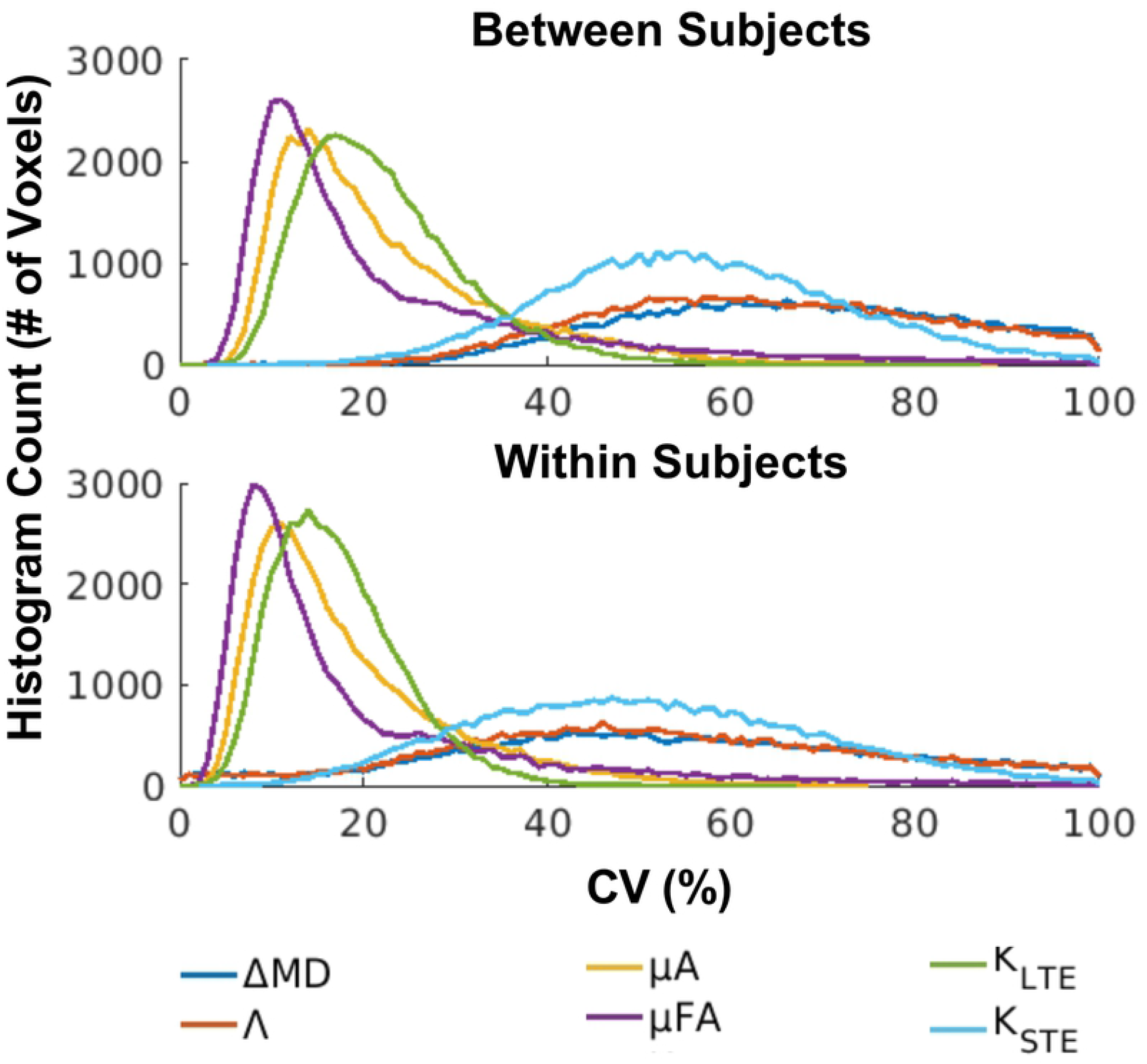
Distribution of between and within subject whole brain voxel-wise CVs for the OGSE and µA metrics.

### Sample sizes and minimum detectable effect

#### Between subjects

The between subject CVs, from the ROI analysis, were used to determine the minimum sample sizes required to detect statistically significant changes of 4, 6, 8, 10, and 12 % between subjects in each metric within each ROI. With a sample size of 12, a minimum change of 4 % could be detected in all ROIs for Λ (**Figure 10**). In comparison, ΔMD required a sample size of 15 to detect a 10 % change in the three larger ROIs (the corpus callosum, hippocampus, and cortex). µA and µFA required sample size of 10 to detect a 6 % change in the three larger ROIs. With a sample size of 16, a 10 % change in K_LTE_ could be detected within all ROIs. K_STE_, on the other hand, required much larger sample sizes (at least 50 subjects per group are required to detect a 12 % change in the cortex).

**Fig 10.**
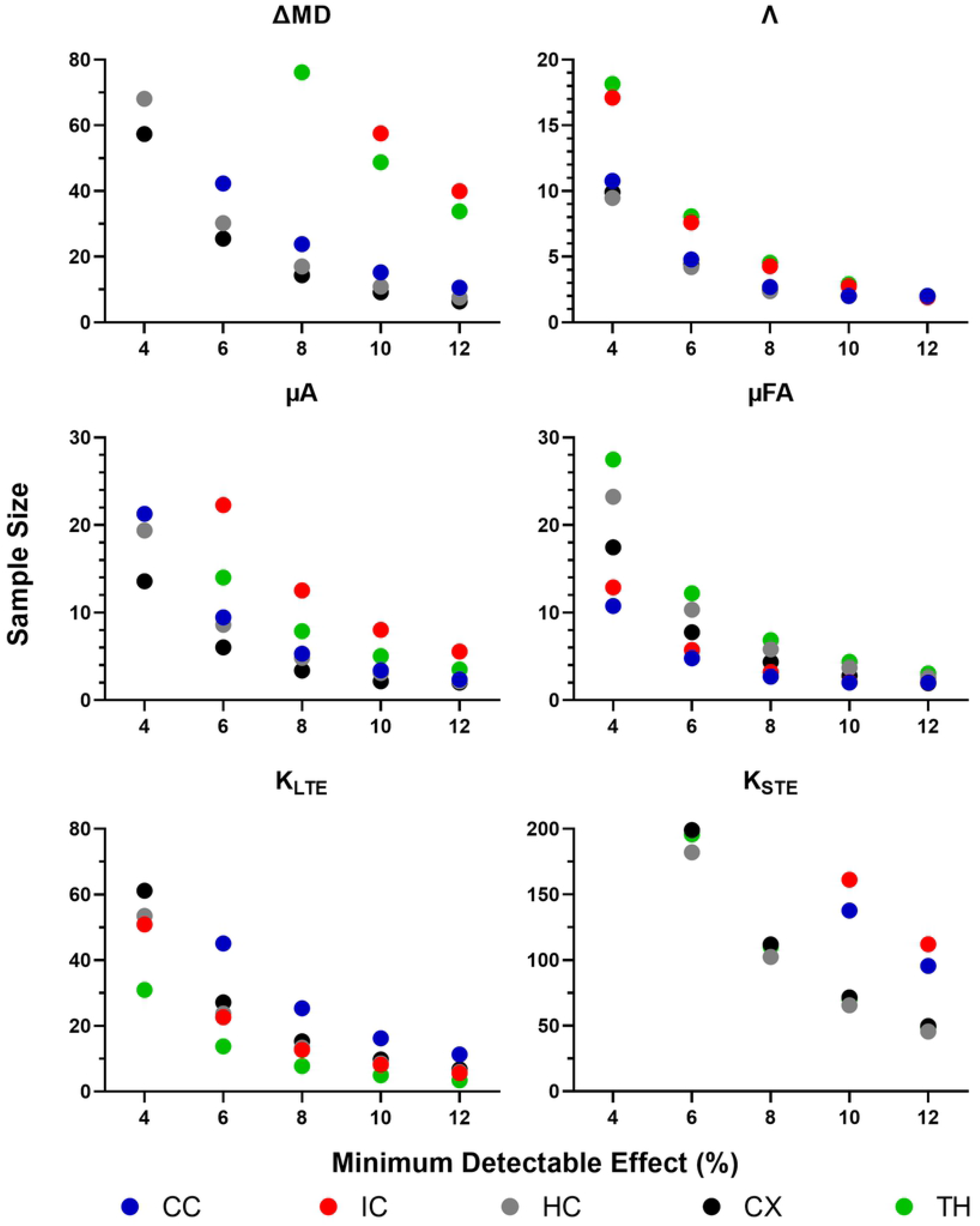
Sample size estimation using a between-subjects approach. Sample sizes required, calculated from ROI-based between-subject CVs, to detect a statistically significant effect within each ROI with a change in each metric of 4, 6, 8, 10, and 12 %. Note that the sample size range varies between plots and sample sizes exceeding the range are not shown. ROIs are abbreviated as follows: CC – corpus callosum; IC – internal capsule; HC – hippocampus; CX – cortex; TH – thalamus.

#### Within subjects

The standard deviations of the differences between test-retest mean values across subjects (assuming paired t-tests) were used to determine the minimum sample sizes required to detect statistically significant changes of 4, 6, 8, 10, and 12 % within subjects in each metric within each ROI.. In the larger ROIs, small changes (6 – 8 %) could be detected in Λ with 10 subjects per group, while ΔMD showed similar trends in the cortex (the largest ROI), but could only detect larger changes (10 -12 %) with the same number of subjects in the corpus callosum and the hippocampus (**Figure 11**). µA was able to detect a minimum change of 4 % in the larger ROIs with 10 subjects per group, while the smaller ROIs required much greater sample sizes. µFA was slightly more robust, being able to detect a 4 % change in the larger ROIs (with 9 subjects per group) and in all ROIs (with 14 subjects per group). K_LTE_ was able to detect moderate changes (8 %) with 12 subjects and smaller changes (6 %) with 15 subjects, whereas K_STE_ required at least 30 subjects to detect larger changes (12 %).

**Fig 11.**
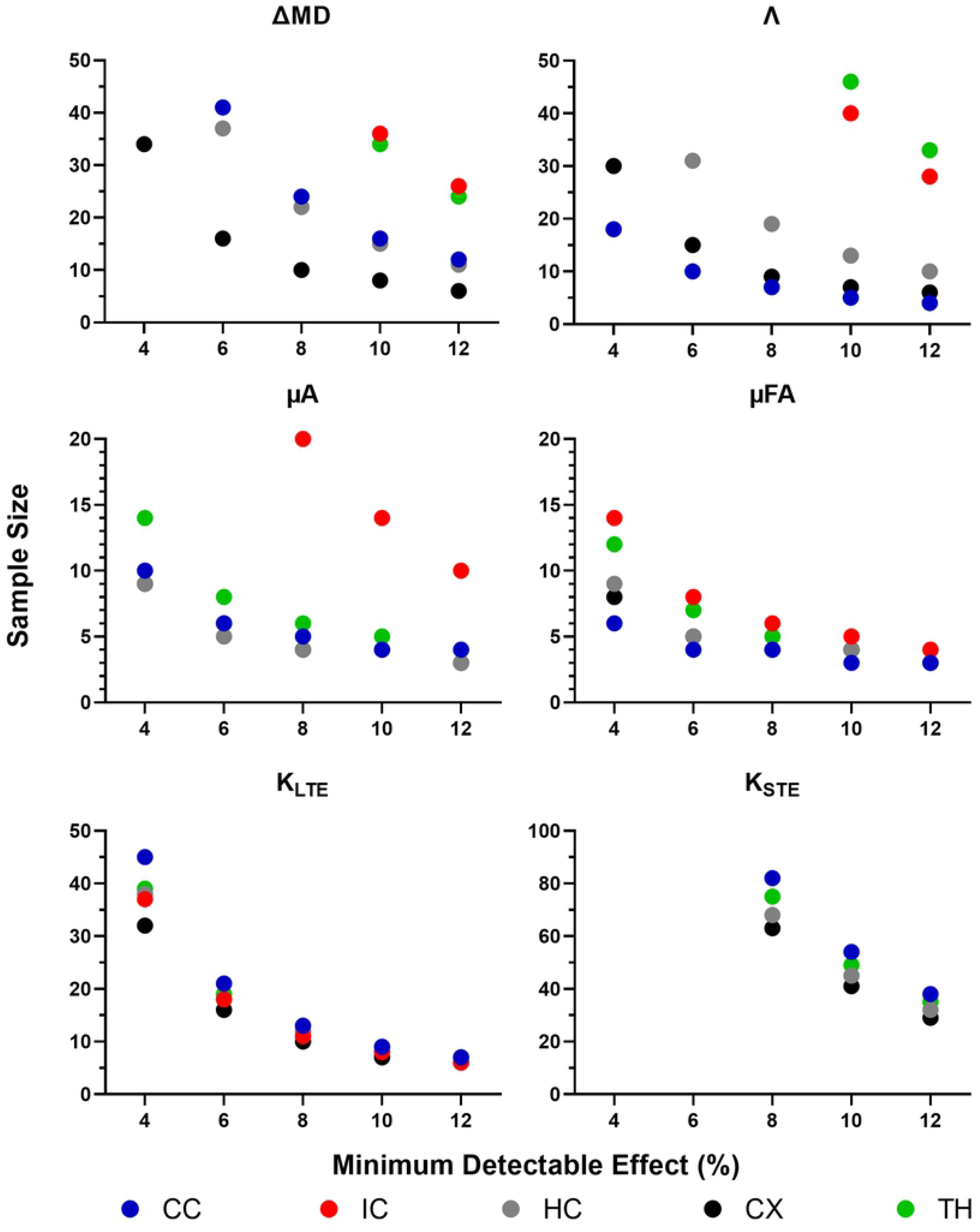
Sample size estimation using a within-subjects approach. Sample sizes required, calculated from the standard deviation of differences between test-retest mean values across subjects (assuming paired t-tests), to detect a statistically significant effect within each ROI with a change in each metric of 4, 6, 8, 10, and 12 %. Note that the sample size range varies between plots and sample sizes exceeding the range are not shown. ROIs are abbreviated as follows: CC – corpus callosum; IC – internal capsule; HC – hippocampus; CX – cortex; TH – thalamus.

## DISCUSSION

This study explored the reproducibility of OGSE and µA metrics at 9.4 Tesla. No biases were found between repeat measurements with either ROI-based or voxel-wise analysis. µA, µFA, and K_LTE_ were shown to be reproducible in both the mean ROI analysis and the whole brain voxel-wise analysis, while ΔMD and Λ were reproducible in only the mean ROI analysis, and reproducibility for K_STE_ could not be established in either the ROI-based or voxel-wise analysis. µA and µFA showed the highest reproducibility of all the metrics and the least dispersion of CVs. The CVs observed for µFA in this work are consistent with CVs reported in a recent study by Arezza *et al*. [18] in human subjects at 3 T, where CVs ranged from 6 – 8 %. Overall, within subject CVs were lower than between subject CVs for both ROI-based and voxel-wise analysis, indicating less variability within subjects on a test-retest basis.

Our ΔMD maps (**Figure 2**) show contrast which is consistent with recent observations in both *in vivo* and *ex vivo* OGSE studies in mouse brains by other groups [23,48,60,61]. Aggarwal *et al*. related the higher OGSE contrast in the dentate gyrus layer of the hippocampal formation to densely packed neurons in the region [48], which simulations have indicated increase the rate of change in MD with frequency [62]. The very low values of ΔMD seen in certain regions of the gray matter are due to partial volume effects from CSF, as CSF exhibits negative values of ΔMD due to flow [12,49]. ΔMD and Λ maps (**Figure 2**) show the same contrast, since the apparent diffusion dispersion rate is directly proportional to ΔMD. This relationship is also reflected in the ΔMD and Λ voxel-wise CV maps (**Figure 8**), which are very similar. While ΔMD requires less scan time than Λ, as it requires only a single OGSE and PGSE acquisition, Λ is expected to be more robust in terms of reproducibility as it includes data from all OGSE acquisitions (as shown in **Figure 3**). This is reflected in our results by the much smaller sample size needed to detect a statistically significant change in Λ, compared to ΔMD (**Figure 10**). In the mean ROI analysis, Λ showed higher between subject reproducibility than ΔMD (**Figure 6**), producing between subject CVs < 5% for all ROIs. An unexpected finding was the comparable within subject CVs for Λ and ΔMD. It should be noted that for the within-subject calculation of CV, the standard deviation was determined from only two data points (the test and retest conditions). As a result, the standard deviation may not accurately represent the spread of data within the population, leading to an unknown bias in the resulting within-subject CV.

In the mean ROI analysis, the size and location of the ROIs influenced the reliability of the measurements. A greater distribution in the mean values for all metrics are observed in the internal capsule and thalamus (**Figure 2**), which are the smallest ROIs analyzed in this study. Similarly, higher CVs and a greater dispersion of CV values are observed in both smaller ROIs (**Figure 4**). This result leads to greater sample sizes being required to detect the same change in the smaller ROIs compared to the larger ROIs (**Figure 10**). Thus, smaller ROIs lead to unreliable measurements due to less averaging and possibly a greater effect from slight registration inaccuracies. Furthermore, both smaller ROIs are positioned in the lower half of the brain, farther away from the surface coil. As CV is inversely proportional to SNR, higher CVs are observed farther away from the surface coil for all metrics (**Figure 8**). To acquire reliable measurements in smaller ROIs and ROIs farther away from the surface coil, greater SNR (corresponding to longer scan time) is required. The use of a volume coil would produce a homogenous SNR (and CV) throughout the whole brain. However, in this study, a surface coil was chosen as it maximized SNR within ROIs in close proximity to the surface coil, such as the cortex and corpus callosum, which are highly reported in rodent neuroimaging studies.

Voxel-wise analysis for specific ROIs (**Supporting Figure 1**) shows that in general, the 3 ROIs shown (the corpus callosum, hippocampus, and cortex) follow the same trends. The corpus callosum shows a slightly lower CV peak than the gray matter regions for the more reproducible metrics (µA, µFA, and K_LTE_). Overall, the within subject CV histograms have peaks at lower values than the between subject CV histograms, indicating less variability on a within subject test-retest basis. This is also noticeable in the between and within subject CV maps (**Figure 8**), with the within subject CV maps showing lower values overall.

One of the main reasons for the lack of reproducibility through voxel-wise analysis of ΔMD and Λ is likely CSF partial voluming. Since voxels with CSF can exhibit negative ΔMD and Λ values, whereas brain tissue shows positive ΔMD values, this leads to very high CVs (CVs > 60) in voxels impacted by CSF contamination, such as in regions with CSF in adjacent slices. This partial volume effect on ΔMD and Λ can be mitigated by using a higher resolution. However, this would also reduce SNR and longer scan times would be required to produce the same image quality. Voxel-wise analysis of ΔMD and Λ (from *in vivo* OGSE data) is not feasible given the resolution and scan time constraints. In contrast, ΔMD and Λ both show good reproducibility in the ROI analysis, where this partial volume effect is mitigated due to averaging. µA, µFA, and K_LTE_ also show greater CVs in regions with CSF, such as the ventricles, arising from the very small values of these metrics in CSF.

As K_STE_ values are intrinsically low in the brain [4,31], higher CVs and greater dispersion of CV values are observed, even in the ROI analysis. Since K_STE_ depends on the variance in mean diffusivity, low K_STE_ values point to a low variance in MD. This indicates similar sized cells across the brain, since a higher variance in cell size would lead to a higher variance in MD. In other words, the volume-weighted variance of cell size is low compared to the mean cell size. It is interesting that although the OGSE metrics (ΔMD and Λ) and K_STE_ all show similar trends in the whole brain voxel-wise analysis (**Figure 9**), the OGSE metrics show greater improvement in CVs with ROI-based analysis than K_STE_. This suggests that averaging (in the ROI-based analysis) does not improve K_STE_ reproducibility, in contrast to the OGSE metrics. Unlike the other metrics explored in this study, K_STE_ shows very low CVs in regions with CSF (**Figure 8**), since K_STE_ values are very high in CSF (**Figure 4**). As the CSF STE signal as a function of b-value decays very rapidly and reaches the noise floor, the fitting detects a false variance (very high K_STE_) if high b-value data is not excluded [4]. The generally low reliability of K_STE_ is likely due to a combination of its low value and the well-known sensitivity of kurtosis fitting to both physiological and thermal noise [63]. Notably, while ostensibly based on kurtosis fitting, µA and µFA do not suffer similar issues because no 2^nd^ order kurtosis fitting is required to estimate these metrics due to term cancellations that occur when the kurtosis difference between LTE and STE is evaluated to estimate these metrics [18]. Despite the low reliability, it is encouraging that the K_STE_ maps (**Figure 2**) exhibit contrast which is comparable to K_STE_ maps shown in a recent *in vivo* rodent study applying correlation tensor imaging (a DDE technique) [38].

Given the current test-retest study design, small changes (< 8 %) can be detected in Λ, µA, and µFA both between and within subjects, with moderate sample sizes of 10 – 15. With all minimum detectable changes explored (**Figure 10** and **Figure 11**), µFA was the most sensitive metric, followed by µA. ΔMD and K_LTE_ can detect such small changes, given a moderate sample size, only within subjects. Between subjects, ΔMD and K_LTE_ can detect moderate changes on the order of 10 %. K_STE_ cannot detect small changes with sample sizes relevant to preclinical neuroimaging studies, unless compromises in scan time or resolution are made to improve SNR compared to the scans performed here.

It should be noted that the findings in this work are specific to the scan parameters used. Diffusion MRI is inherently a low SNR technique and high b-value acquisitions (from the µA protocol) and high oscillating gradient frequency acquisitions (from the OGSE protocol) result in even lower SNR. To acquire sufficient SNR, the voxel size was adjusted, with slice thickness set to 500 µm. Acquiring a greater SNR than the present study would provide more reliable measurements. Since our metrics are greatly impacted by partial volume effects (mostly from CSF), a higher resolution would provide more accurate and reproducible measurements.However, acquiring a higher SNR with higher resolution would require much greater scan time, which is not feasible for longitudinal *in vivo* neuroimaging studies, which are essential to characterize the progression of disease and injury recovery. Furthermore, a single channel transceive surface coil was used in this study and scan acceleration with parallel imaging was not possible. An option for obtaining more reliable ΔMD measures is to acquire only one PGSE and one OGSE scan, utilizing the same scan time of 45 minutes for the multifrequency OGSE protocol in this study. Thus, greater SNR and/or resolution can be achieved with more averaging. However, in doing so, one would lose the potential additional insight into microstructure organization and tissue integrity that multiple frequency analysis can provide if, for example, the f^0.5^ power law scaling of MD changes in certain pathologies.

## CONCLUSION

In conclusion, we have investigated the reproducibility of OGSE and µA metrics in a rodent model at an ultra-high field strength. We have shown that the µA, µFA, and K_LTE_ metrics (from the µA protocol) are reproducible in both ROI-based and voxel-wise analysis, while the ΔMD and Λ metrics (from the OGSE protocol) are only reproducible in ROI-based analysis. Λ, µA, and µFA may provide sensitivity to subtle microstructural changes (4 - 8 %) with feasible sample sizes (10 – 15). This work will provide insight into experiment design and sample size estimation for future longitudinal *in vivo* OGSE and µA microstructural dMRI studies at 9.4 T.

## SUPPORTING INFORMATION

**S1 Fig. Distribution of voxel-wise between and within subject CVs within each ROI**.

## AUTHOR CONTRIBUTIONS

**Conceptualization:** Naila Rahman, Arthur Brown, Corey A. Baron

**Data Curation:** Naila Rahman, Kathy Xu, Corey A. Baron

**Formal Analysis:** Naila Rahman, Mohammad Omer

**Funding:** Corey A. Baron

**Investigation:** Naila Rahman

**Methodology:** Naila Rahman, Kathy Xu, Arthur Brown, Corey A. Baron

**Project Administration:** Naila Rahman

**Resources:** Kathy Xu, Matthew D. Budde, Arthur Brown, Corey A. Baron

**Software:** Naila Rahman, Matthew D. Budde, Corey A. Baron

**Supervision:** Arthur Brown, Corey A. Baron

**Validation:** Naila Rahman, Corey A. Baron

**Visualization:** Naila Rahman

**Writing – original draft:** Naila Rahman

**Writing – review & editing:** Naila Rahman, Kathy Xu, Mohammad Omer, Matthew D. Budde, Arthur Brown, Corey A. Baron

## Notes

### Competing Interest Statement

The authors have declared no competing interest.

